# From presence/absence to reliable prey proportions: A field test of dietary DNA for characterizing seabird diets

**DOI:** 10.1101/2024.03.22.586275

**Authors:** Gemma V Clucas, Andrew Stillman, Elizabeth C Craig

## Abstract

Climate change and human impacts are causing rapid shifts in species’ distributions and abundance, potentially disrupting predator-prey relationships. Monitoring animal diets can elucidate these relationships and potentially provide a mechanistic understanding of population declines, but efficient methods to monitor diets are needed. Dietary DNA can be used to gather data on species’ diets with high-taxonomic resolution, but the question of whether it can provide quantitative information on animal diets has so far limited its applications. Here we show that dietary DNA can efficiently provide quantitative information on the relative proportions of fish in the diets of piscivorous seabirds. Over three breeding seasons, we observed common tern chick feeding events and collected fecal samples from chicks. We compared the frequency of occurrence (FOO) and relative read abundance (RRA) of fish prey in the fecal samples to the relative biomass recorded through visual observations and found a high correlation (R = 0.94) between RRA and relative biomass at the colony level. We also found that RRA outperformed FOO in capturing interannual changes in relative prey biomass, and that FOO systematically over-estimated the relative importance across all prey categories. The high taxonomic resolution provided by dietary DNA and the high correlation between RRA and relative biomass we found here suggest that dietary DNA can be used as a quantitative metric to monitor seabird diets. This has important applications for studying the impact of changing forage fish availability on seabird populations, but also for harnessing seabirds as sentinels of ecosystem health. Since quantitative diet data can be collected highly efficiently using dietary DNA, this method could provide key information about food web dynamics and forage fish availability as we move towards ecosystem-based fisheries management.

## 1. Introduction

Species extinctions and population declines continue at alarming rates (Barnosky et al., 2011; Ceballos et al., 2017; Dirzo et al., 2014; Pimm et al., 2014; Rosenberg et al., 2019), and addressing these declines requires a mechanistic understanding of the complex drivers and environmental conditions that impact populations. Monitoring animal diets can provide a critical window into these mechanisms. Diet may be highly influenced by environmental conditions and resource availability, and diet in turn can impact an individual’s survival, reproduction, and fitness (Spiller and Dettmers, 2019). Dietary DNA has become an increasingly powerful and popular tool to study these links over the last 15 years due to the ease and taxonomic resolution afforded by dietary DNA over traditional diet monitoring techniques (de Sousa et al., 2019; Pompanon et al., 2012; Valentini et al., 2009). Frequently, fecal samples are used in dietary DNA studies as they can be collected non-invasively, although the method can also be used to identify species from gut contents or regurgitated material. To identify prey species, dietary DNA studies may employ one of three techniques: 1) using targeted PCR to detect prey species of interest through the use of highly specific PCR primers; 2) using a metabarcoding approach, whereby barcoding genes are amplified and sequenced from a broad range of taxa using degenerative or universal primers; and 3) using a metagenomic approach, whereby all DNA molecules contained in the sampled material are sequenced. Fecal metabarcoding is by far the most popular of these methods because of its relative efficiency (vs metagenomic approaches) and its ability to identify multiple species simultaneously (vs targeted PCR), making it a powerful tool to increase our mechanistic understanding of conservation challenges in the Anthropocene.

A major limitation in the use of dietary DNA in conservation and management is the notion that it cannot produce quantitative estimates of animal diets (Hoenig et al., 2022). That is, the proportion of sequence reads (relative read abundance or RRA) recovered from each prey species does not reflect prey biomass consumed. Indeed, there are many factors that can distort RRA in dietary DNA studies, including prey-specific DNA content and digestibility (Thomas et al., 2014), secondary consumption (Bowser et al., 2013), contamination from the environment or in the lab, PCR stochasticity, preferential amplification and sequencing of certain prey sequences over others, and bioinformatic biases (see Alberdi et al., 2019 for a full review). Captive feeding studies and metabarcoding of tissue mixtures have shown variable results (reviewed in Deagle et al., 2019) with RRA often showing a correlation with biomass but with low accuracy. Therefore, dietary DNA results are often converted to presence/absence data as a conservative alternative to RRA. However, Deagle et al. (2019) showed through simulations that converting sequence counts to presence/absence data can over-estimate the importance of rare diet items because they are counted equally alongside items consumed in greater proportions. Secondary consumption and contamination can also have a greater impact on presence/absence results. Therefore, RRA may give a more accurate representation of population-level diet compared to presence/absence in many circumstances (Deagle et al., 2019). At present, however, few studies have validated this method in wild populations (Verkuil et al., 2020), and more system-specific validation studies are needed before dietary DNA can reach its full potential as an effective tool for conservation.

Seabirds are one of the most threatened groups of vertebrates (Croxall et al., 2012). Monitored populations have declined by 70% between 1950 - 2010 (Paleczny et al., 2015), and threats from interactions with fisheries and climate change continue to raise conservation concerns for these species (Dias et al., 2019; Paleczny et al., 2015; Phillips et al., 2022). Fisheries-induced mortality may occur directly through bycatch, particularly in long-line and trawl fisheries, or indirectly, through competition for prey resources (Anderson et al., 2011; Phillips et al., 2022). Climate-linked seabird declines are often indirect and the result of changing prey resources. For example, climate-induced shifts in prey availability and over-fishing of key prey species has caused declining breeding success of Atlantic puffins in the North Atlantic (Durant et al., 2003; Fayet et al., 2021; Kress et al., 2016; Scopel et al., 2019). Since seabirds tend to forage across large geographic areas, often nest in large colonies, and feed primarily on forage fish and other hard-to-monitor prey species, they have long been recognized as sentinels of marine ecosystem health (Ainley et al., 1996; Cairns, 1987; Furness and Camphuysen, 1997; Piatt et al., 2007), with their diets providing quantitative estimates of the availability of forage fish and squid (Diamond and Devlin, 2003; Montevecchi, 1993; Montevecchi and Myers, 1995; Scopel et al., 2017; Velarde et al., 2019, 1994). However, traditional methods to monitor their diets through stomach content analysis, stable isotopes, or visual observations of chick feeding events frequently lack taxonomic resolution due to the difficulty in identifying prey to species level. This lack of taxonomic specificity contributes to uncertainty in how preyscapes critical to seabird survival will shift under future climate scenarios. Traditional methods can also be invasive or restricted to bill-loading birds. As an alternative, tracking changes in diet through quantitative dietary DNA could greatly increase our ability to monitor real-time impacts on seabird populations and proactively address fisheries and climate-driven threats.

Our objectives in this study were to (1) determine the extent to which dietary DNA can estimate fish prey proportions in a wild seabird population and (2) investigate the effects of sample size on these estimates. Our focal system was the common tern (*Sterna hirundo*) nesting in the Gulf of Maine, where visual observations have historically been used to monitor prey fed to chicks by adult terns. We compared chick dietary DNA to these visual observations and focused exclusively on fish, excluding the analysis of invertebrate prey, since fish are the primary food source for these birds (Arnold et al., 2020). To validate dietary DNA against visual observations, we calculated four metrics commonly used in diet studies:

1) Relative count: the relative proportion of prey taxa based on counts from visual observations.
2) Relative biomass: the relative proportion of prey taxa based on estimated prey biomass from visual observations.
3) FOO (frequency of occurrence): the proportion of samples in which prey taxa are present in the dietary DNA (i.e. presence/absence of prey species).
4) RRA: the relative proportion of DNA sequences of prey taxa in the dietary DNA.

Relative prey counts are often used to estimate prey proportions, but relative biomass estimates provide a more informative metric to characterize diets when data on prey size are available. Thus, we focused our inference on assessing whether dietary DNA could capture the relative biomass of prey. We also simulated an annual monitoring scheme to investigate the effects of sample size on the ability of dietary DNA to match estimates from visual observations. If dietary DNA can match the inferential capabilities of traditional methods such as visual observations, this approach opens doors to track real-time changes in prey availability and more efficiently address the impacts of fisheries management and climate change on prey resources critical to seabird populations.

## 2. Methods

### 2.1 Study site and species

This study was conducted at a mixed-species tern breeding colony on White and Seavey islands (42°58’ N, 70°37’ W) located in the Isles of Shoals archipelago, New Hampshire, USA, within the Gulf of Maine. These islands are New Hampshire’s primary breeding colony for common, roseate (*S. dougallii*), and Arctic terns (*S. paradisaea*), and the colony is a key conservation area with regional significance. Restoration, management, and research activities have been conducted at this site since 1997 under the auspices of the New Hampshire Fish and Game Department. Data presented here were collected in June and July of 2017, 2018, and 2019 during routine monitoring activities. In the years of this study, the colony annually supported 2000–3000 pairs of common terns, 60–90 pairs of roseate terns, and 1–2 pairs of arctic terns.

### 2.2 Visual observations

Common terns provision their chicks with whole prey items, typically fish of 0 to 4 years of age and up to 15cm in length (Arnold et al., 2020). Visual observations of chick provisioning were conducted in June and July of each year, either in-person with binoculars from observation blinds or remotely using video footage recorded from an AXIS P5635-E Mk II PTZ Network Camera. Observations occurred between the hours of 05:30 and 21:30. Visual provisioning data were collected in 2017 from video recordings at 13 nests for a total of 160 nest hours, in 2018 from video recordings at 14 nests for a total of 167 nest hours, and in 2019 from both video recordings and in-person observations at 22 nests for a total of 310 nest hours.

For each provisioning event, prey items were visually identified to the highest possible taxonomic resolution. While tern diets included both fish (described below) and invertebrate prey (primarily euphausiids, insects, and squid), invertebrates were not within the scope of this study and excluded from further analysis. Prey fish were grouped into six categories:

(1) Hake (likely *Urophycis chuss, U. tennuis,* and *Merluccius bilinearis*).
(2) Herring (likely *Clupea harengus* and *Alosa spp*.).
(3) Sand lance (*Ammodytes* spp.).
(4) Atlantic butterfish (*Peprilus triacanthus*).
(5) Other identifiable fish.
(6) Unknown fish (including fish that were unidentifiable due to the speed of provisioning, obscured view of identifying features, or lack of confidence on behalf of the observer).

Prey size was quantified by visually estimating the length of each prey item relative to that of the adult tern’s bill to the nearest 0.5 bill lengths, then multiplying by average adult bill length at our study site (3.65 cm). The biomass of each fish was calculated by inserting our estimate of fish length into length-weight relationships from the literature (Schneider et al., 2000; Wigley et al., 2003; Winters, 1989). The length-weight equation for Atlantic herring was used for all fish in the herring category, and the equation for white hake was used for all fish in the hake category. For identifiable fish species with no published length-weight relationships, we substituted equations of morphologically similar species for which equations were available. The equation for white hake was also used for fish in the unknown fish category, as white hake has a narrow, fusiform shape, typical of the morphology of many unidentified fish in our study. Provisioning events were further categorized into successful or failed feedings depending on whether the provisioned prey was successfully consumed by the chick. Failed feedings, where prey was either stolen or discarded, were excluded from further analysis for this study.

### 2.3 Fecal sample collection

We collected fecal samples from common tern chicks (≤ 2 weeks old) if they defecated during handling as part of standard monitoring procedures. Fecal samples were transferred to 2ml tubes pre-filled with 1 ml of DNA/RNA Shield preservation buffer (Zymo Research, Irving, CA) using single-use, individually-wrapped spatulas. Samples were only collected from “clean” surfaces, such as pre-cleaned weighing scales, bird bags, clothing, or hands. To monitor for DNA contamination from these surfaces, we collected negative control samples (hereafter “field blanks”) from all surfaces by dipping a spatula into the preservation buffer to dampen it, wiping it on the surface to be sampled two or three times to simulate picking up a sample, and stirring it back into the preservation buffer as if depositing a sample into the tube. We treated field blanks equally to fecal samples for subsequent metabarcoding steps. Samples were stored at ambient temperature, out of direct sunlight, for up to 1 year until DNA extractions took place.

### 2.4 DNA extractions and metabarcoding

DNA extractions were performed using Zymo Research Quick-DNA Fecal/Soil Miniprep 96 kits. We found that the 96-well bead bashing plates supplied in the kit would leak during beating, so we transferred each sample and its preservation buffer to an individual bead bashing tube to avoid cross-contamination. We bead beat samples on a benchtop vortex set at maximum speed for 20 minutes. The rest of the DNA extraction was performed following the manufacturer’s instructions after transferring the centrifuged supernatant from the bead bashing tubes into the kit’s 96-well filter plates. We eluted the DNA into a final volume of 60 µl of elution buffer. Extractions were performed alongside other tern fecal sample extractions, with samples mixed among plates to minimize batch effects. We included eight extraction blanks (no template controls) in each plate to monitor for cross-contamination during extractions and performed all DNA extractions and PCR set-ups in a dedicated clean lab.

Dietary metabarcoding was performed with a 2-step PCR approach. In the first step, the 12S rRNA gene was amplified using MiFish primers (Miya et al., 2015; underlined below) to which we had added TruSeq tails (primer sequences were thus MiFish-U-F-TruSeq: 5’-ACACTCTTTCCCTACACGACGCTCTTCCGATCT<u>GTCGGTAAAACTCGTGCCAGC</u>-3’; and MiFish-U-R-Truseq: 5’-TGACTGGAGTTCAGACGTGTGCTCTTCCGATCT<u>CATAGTGGGGTATCTAATCCCAGTTTG</u>-3’). We included a PCR blank (no template control) with every batch of PCRs to monitor for contamination during PCR setup. A mock community (positive control), including DNA extracted directly from fish samples, was also included and added to the plate last. The PCR reaction included 6 μL of KAPA HiFi HotStart ReadyMix 2X (KAPA Biosystems, Wilmington, Massachusetts), 0.7 μL of each of the forward and reverse primers at 5 μM concentration, and 4.6 μL of fecal DNA (or molecular grade water for the PCR blanks). For the mock community, we reduced the amount of template DNA to 1 μL and added 3.6 μL of molecular grade water. Thermocycling conditions were: 95 °C for 3 mins, 35 cycles of 98 °C for 30 secs, 65 °C for 30 secs, 72 °C for 30 secs, followed by a 5-minute extension at 72 °C. PCR products were visualized using gel electrophoresis to ensure that all blanks were clear. PCRs were performed in duplicate to minimize the effects of PCR stochasticity, with the products from two replicates being combined and diluted depending on the strength of their bands before the second stage PCR.

The second stage PCR used the diluted product from the first stage as template and added the flow cell binding sites and sequencing primer binding sites. The primer sequences were: forward, 5’-AATGATACGGCGACCACCGAGATCTACAXXXXXXXXACACTCTTTCCCTACACGAC-3’ and reverse, 5’- CAAGCAGAAGACGGCATACGAGATXXXXXXXXGTGACTGGAGTTCAGACGTGT-3’. The octo-X sites represent the i7 and i5 indexes used to identify samples. We used unique dual indexes, such that any reads that suffered from tag-jumping would be eliminated during de-multiplexing. The PCR reaction used 6 μL of KAPA HiFi HotStart ReadyMix 2X, 0.7 μL of each of the forward and reverse primers at 5 μM concentration, 1 µL of template and 3.6 µl of molecular grade water. The thermal cycling profile was 94 °C for 3 mins, 15 cycles of 94 °C for 20 secs, 72 °C for 15 secs, and a final extension of 7 mins at 72 °C. Sequencing was performed on an Illumina NovaSeq 6000 using an SP flow cell with 250bp paired-end reads at the University of New Hampshire’s Hubbard Center for Genome Studies. Samples were loaded alongside other projects using a small percentage of a lane.

### 2.5 Bioinformatics

Bioinformatics were performed using Qiime2 v2021.4 (Bolyen et al., 2019). First we trimmed forward and reverse primers using the cutadapt plugin (Martin, 2011). We then denoised and merged paired-end reads using the DADA2 plugin (Callahan et al., 2016), truncating the forward and reverse reads to 133 and 138 bp, respectively, and specifying a minimum overlap of 50 bp between them. After denoising, we merged all the data across the different sequencing plates and then assigned taxonomy to the sequences using an iterative BLAST method and a custom reference database. To create the reference database, we used the RESCRIPt plugin (Robeson et al., 2021) to download any 12S or mitochondrial genomes from GenBank that originated from fish or birds that were studied in our lab. Downloaded database sequences were cleaned and dereplicated using RESCRIPt default parameters and finally, a human mitochondrial genome was added to the database, as this is a common source of contamination. The iterative BLAST method then took each representative sequence from our samples and blasted it 80 times against the reference database, increasing the percent identity incrementally from 70 – 100 %, thus circumventing the limitation of the BLAST method, which keeps only the first hit that meets the search criteria, rather than the best hit. The script for the iterative BLAST method is available from https://bitbucket.org/dwthomas/qiime2_tools/src/master/mktaxa.py. We chose this method to assign taxonomy rather than training a Naïve Bayes classifier, as this method identified all species in our mock community correctly, while a trained classifier mis-identified one species (Atlantic butterfish). Next, we filtered out unassigned reads, reads originating from human contamination, and reads originating from the birds themselves.

Alpha-rarefaction curves showed that 400 reads were necessary to capture the fish diversity in the fecal samples. We therefore normalized all samples to a depth of 400 reads and dropped any that had fewer. We then manually checked all species assignments by blasting representative sequences against the full GenBank database and checked that the fishes’ ranges included the Gulf of Maine using FishBase (www.fishbase.se). Where multiple species occurred in the Gulf of Maine and each had a 98% match or higher with our reference sequence, such as sand lances (*Ammodytes dubius* and *Ammodytes americanus*) and river herrings (*Alosa aestivalis, Alosa pseudoharengus,* and *Alosa sapidissima*) we did not attempt to make species assignments and instead used genus-level assignments.

### 2.6 Comparisons with visual observations

We summarized results from visual observations and dietary DNA using four separate metrics which describe diet composition: relative counts, relative biomass, FOO, and RRA. Using data from visual observations, we calculated relative counts by dividing the number of observations in each prey group by the total number of prey observations. Similarly, we calculated relative biomass by dividing the observed biomass in each group (i.e., number of prey × mass) by the total biomass across observations. Next, we used dietary DNA results to calculate FOO and RRA. Metrics based on dietary DNA had a higher taxonomic resolution than metrics based on visual observations, so we grouped summaries into the following categories to match visual observations: 1) hake, which included red hake, white hake, and silver hake; 2) herring, which included Atlantic herring and river herrings; 3) sand lance; 4) Atlantic butterfish; and 5) other, which included all other species detected in the fecal samples. Unlike the visual observations, there was no “unknown fish” category. We calculated all four metrics separately for the three sampling years and for all years combined. To facilitate comparisons, we omiwed “other” and “unknown fish” categories from statistical tests, and we completed analyses using R version 4.2.2 (R Core Team, 2022) aided by the package “FSA” (Ogle et al., 2023).

We used each prey category-year combination as the sampling unit for statistical tests, resulting in a sample size for comparisons of n = 48. First, we calculated the Pearson correlation coefficient for each pair of metrics. Next, we calculated the mean absolute error (MAE) and root mean squared error (RMSE) to measure differences between each pair of metrics while accounting for the magnitude of differences. RMSE is based on the average of squared differences between two metrics, and comparisons based on RMSE tend to give a higher weight to large errors. We compared MAE and RMSE between the four dietary metrics using a Kruskal-Wallis rank sum test followed by a post-hoc Dunn test.

We also compared the ability of the metrics to track interannual changes in prey consumption. To accomplish this, we calculated the proportional change between years for each metric using the equation

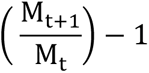

Where **M_t_** is the value of a diet metric in an initial year, and **M_t+1_** is the value of the same metric in the following year. The results of this equation ranged from-0.90 to 7.80 and can be interpreted as the change between consecutive years, expressed as a proportion of the initial value. We then calculated the Pearson correlation coefficient for interannual changes detected by each pair of metrics (herea{er “interannual correlation”).

Finally, we used a simulation study to test how the relationship between visual observations (specifically relative biomass) and dietary DNA (specifically RRA) changed based on the number of fecal samples collected per year. This approach can (a) investigate whether additional fecal sampling could further improve the correspondence between relative biomass and RRA in this study system, and (b) inform sampling design considerations for future diet monitoring efforts using dietary DNA. To accomplish these objectives, we started by sequentially subsampling the dietary DNA dataset under a simulated sampling design ranging from 10–40 fecal samples/year. We sampled without replacement for 2018 and 2019. For 2017 (n = 30) and simulations of > 30 samples/year, we used the full set of 30 samples and then resampled from 2017 to reach the desired number. We repeated this procedure for 500 iterations given each sampling design and calculated RRA for each species category-year at every iteration, resulting in a dataset of 500 calculations for each sampling design ranging from 10 to 40 samples/year. Using the relative biomass for each species category-year as our observational “truth”, we then calculated MAE, RMSE, and interannual correlation between relative biomass and RRA for each simulated dataset and iteration.

## 3. Results

### 3.1 Visual observations

We recorded 967 successful feeding events over 637 nest observation hours during the three years of this study (Table 1). Based on visual observations, the most frequently observed prey categories were hake, herring, and unknown fish (Figure 1). A large proportion of visual observations fell into the unknown fish category (Figure 1) as the size, speed, or angle at which fish were provisioned did not allow for a positive species identity to be determined by eye in nearly a third of all provisioning observations (N=297). These unknown fish made up a large proportion by count, but a smaller proportion by biomass, as unidentified fish were smaller on average (5.8 ± 2.0 cm for unidentified fish compared to 6.9 ± 2.4 cm for fish with positive prey IDs). Prey species within the “other identifiable fish” category included, in order of relative frequency, mummichogs (*Fundulus heteroclitus*), cunner (*Tautogolabrus adspersus*), fourbeard rockling (*Enchelyopus cimbrius*), other gadid species (likely haddock [*Melanogrammus aeglefinus*], pollock [*Pollachius virens*], and Atlantic cod [*Gadus morhua*]), lumpfish (*Cyclopterus lumpus*), Atlantic silverside (*Menidia menidia*), white sucker (*Catostomus commersonii*), and Northern pipefish (*Syngnathus fuscus*). Hake had the highest relative count in the diet for all years combined, but the higher mass of herring (average = 4.2 ± 3.6 g) compared to hake (1.4 ± 1.3 g) showed that herring contributed more to the total biomass of nestling diets (Figure 1).

**Figure 1.**
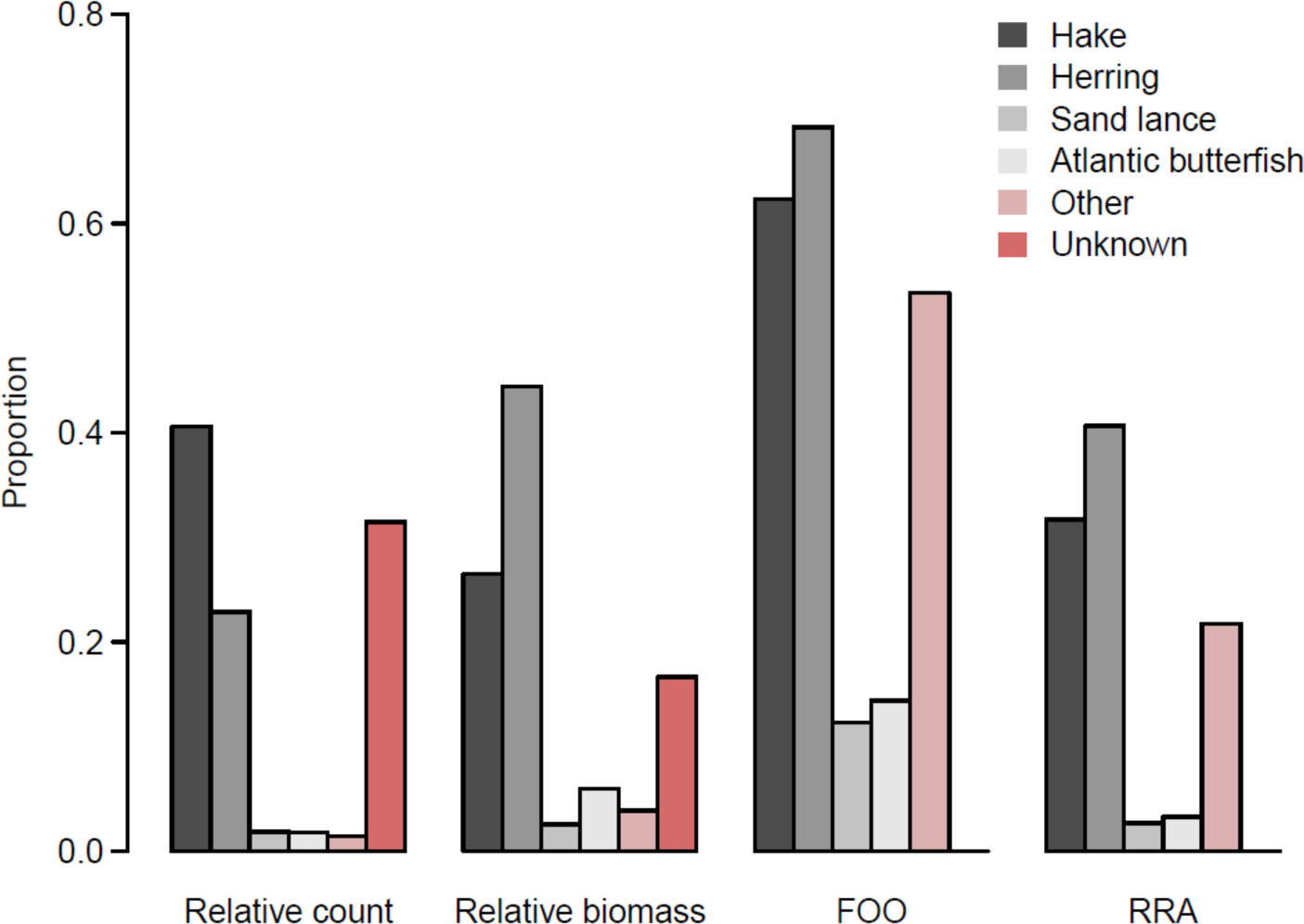
Diet characterization for wild common tern (*Sterna hirundo*) nestlings observed at a Gulf of Maine breeding colony from 2017-2019. Diet metrics summarized from visual observations included relative count and relative biomass. Diet metrics summarized from dietary DNA include frequency of occurrence (FOO) and relative read abundance (RRA).

**Table 1.** The number of visual observations and dietary DNA samples used in each year to study the diets of common terns.

| Year | Number of feeding events observed | Fecal samples collected | Fecal samples included in final analysis |
| --- | --- | --- | --- |
| 2017 | 113 | 40 | 30 |
| 2018 | 163 | 55 | 42 |
| 2019 | 690 | 80 | 74 |

### 3.2 Dietary DNA from fecal samples

Overall, we had a high sequencing success rate, allowing us to use 81% of samples in our final analysis after all filtering steps (Table 1). Samples, blanks, and positive controls yielded nearly 63 million sequences in total, with a mean of 248,725 reads per sample and 219,574 reads per blank. After adapter removal, denoising, and chimera removal, 73% of reads were retained in the samples and 0.03% were retained in the blanks on average. After removing unassigned, human, and avian sequences and normalizing the depth of all samples to 400 reads, all field-, extraction-, and PCR blanks were dropped because they had fewer than 400 reads remaining. Contamination from sample collection surfaces in the field and from cross-contamination in the lab thus appears to have been negligible. Similar to visual observations, dietary DNA approaches revealed the most prevalent prey categories to be hake and herring across all years.

### 3.3 Comparison of visual observations and fecal metabarcoding

We recorded 12 fish prey groups (including eight within the “other identifiable fish” category) in our visual observations compared with 19 in our fecal samples. This was largely due to the greater taxonomic resolution afforded by our dietary DNA data. For example, we could differentiate Atlantic herring from river herrings, and gadid species from one another, including hakes, haddock, and Atlantic tomcod, all of which resemble one another as juveniles. Logistical constraints on visual observations resulted in a large proportion of prey items categorized as “unknown”, while there were no unidentifiable prey in the dietary DNA (Figure 1). As a result, dietary DNA summaries estimated a higher proportion of prey categorized as “other” compared to visual observations, reflecting the high taxonomic resolution of DNA approaches.

Pairwise correlations between each of the dietary metrics examined in this study are summarized in Table 2. Summing across all years of the study, the highest correlation with relative biomass was RRA (r = 0.94; Table 2, Figure 2a). In fact, the strength of this correlation was far higher than the correlation between relative biomass and relative counts and similar to the correlation between RRA and FOO (Table 2). FOO also showed a strong correlation to relative biomass (r = 0.92; Table 2). However, FOO systematically over-estimated the relative importance of nearly all prey categories in all years (Figure 2b) because FOO can range from 0 to 1 in each prey category, while all other metrics (RRA, relative counts, and relative biomass) sum to 1 across all prey categories combined.

**Figure 2.**
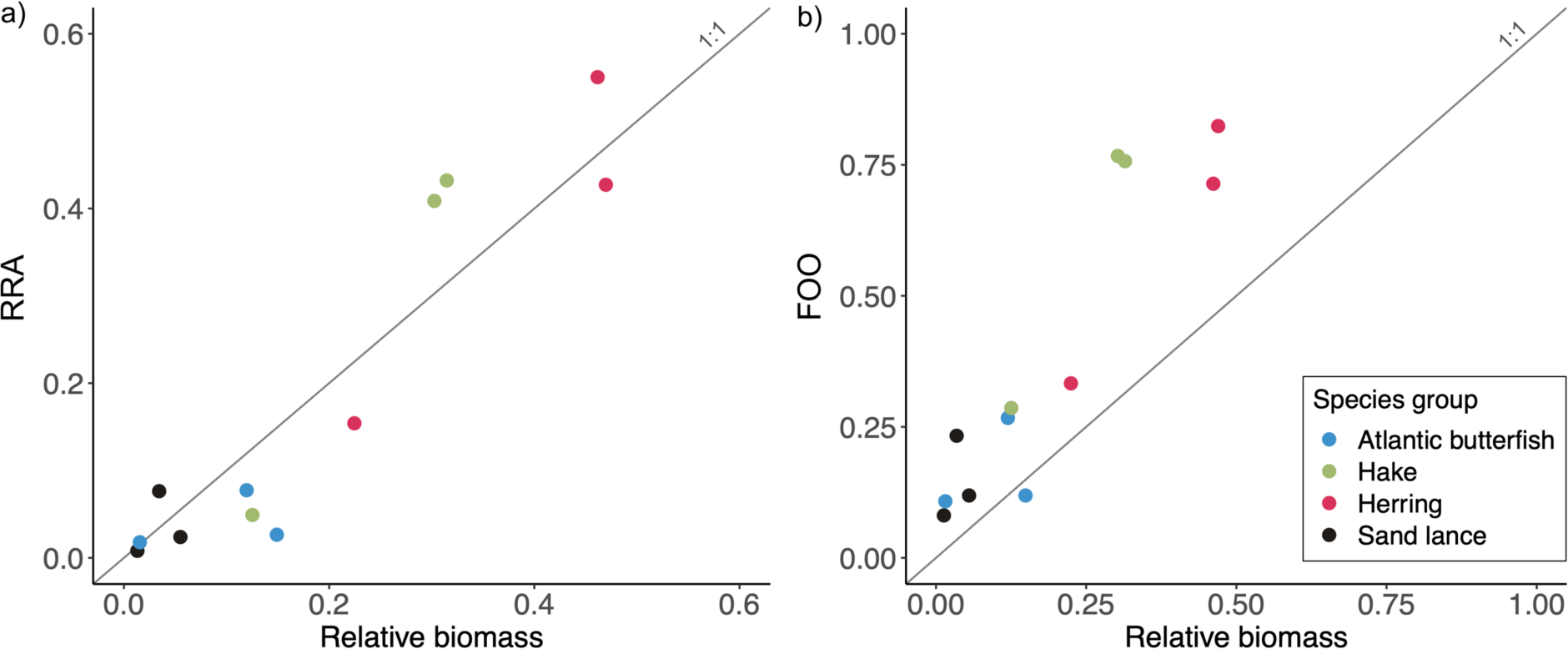
a) The high correlation between RRA and relative biomass (r = 0.94) falls close to a 1:1 relationship. b) FOO also shows a high, positive correlation (r = 0.92), but tends to overestimate the relative importance of most prey groups relative to their biomass.

**Table 2.** Correlation coefficients and errors among four diet metrics characterizing the diets nestling common terns in the wild. Relative counts and relative biomass come from visual observations, and relative read abundance (RRA) <u>and frequency of occurrence (FOO) come from dietary DNA.</u>.

|  | Correlation | MAE | RMSE |
| --- | --- | --- | --- |
| Relative biomass vs RRA | 0.94 | 0.062 | 0.074 |
| Relative biomass vs FOO | 0.92 | 0.199 | 0.245 |
| Relative biomass vs relative counts | 0.66 | 0.104 | 0.133 |
| RRA vs relative counts | 0.92 | 0.087 | 0.139 |
| RRA vs FOO | 0.96 | 0.196 | 0.223 |
| FOO vs relative counts | 0.81 | 0.229 | 0.284 |

MAE and RMSE measurements for each pair of metrics showed strong differences between comparisons (Kruskal-Wallis χ^2^ = 14.5, p = 0.002). MAE and RMSE error rates were lowest when comparing RRA to relative biomass and highest when comparing FOO to relative counts and relative biomass (Table 2). Notably, error rates from the RRA vs. relative biomass comparison were even lower than comparisons based on the same data collection techniques, such as relative biomass vs. relative counts (both summarized from visual diet observations) and RRA vs. FOO. Post-hoc tests showed strong evidence that RRA provided a better match to relative biomass (Dunn test Z = 2.5, p = 0.04) and relative counts (Dunn test Z = 2.9, p = 0.021) compared to FOO. When examining year-to-year changes in diet metrics across four focal prey categories (Figure 3), the interannual correlation between dietary DNA metrics and relative biomass was strong for both RRA (r = 0.77) and FOO (r = 0.74). Interannual correlations were substantially weaker when comparing relative counts to RRA (r = 0.14) and FOO (r = 0.11), highlighting the usefulness of biomass-based measurements over relative counts for detecting and tracking shifts in diets over time.

**Figure 3.**
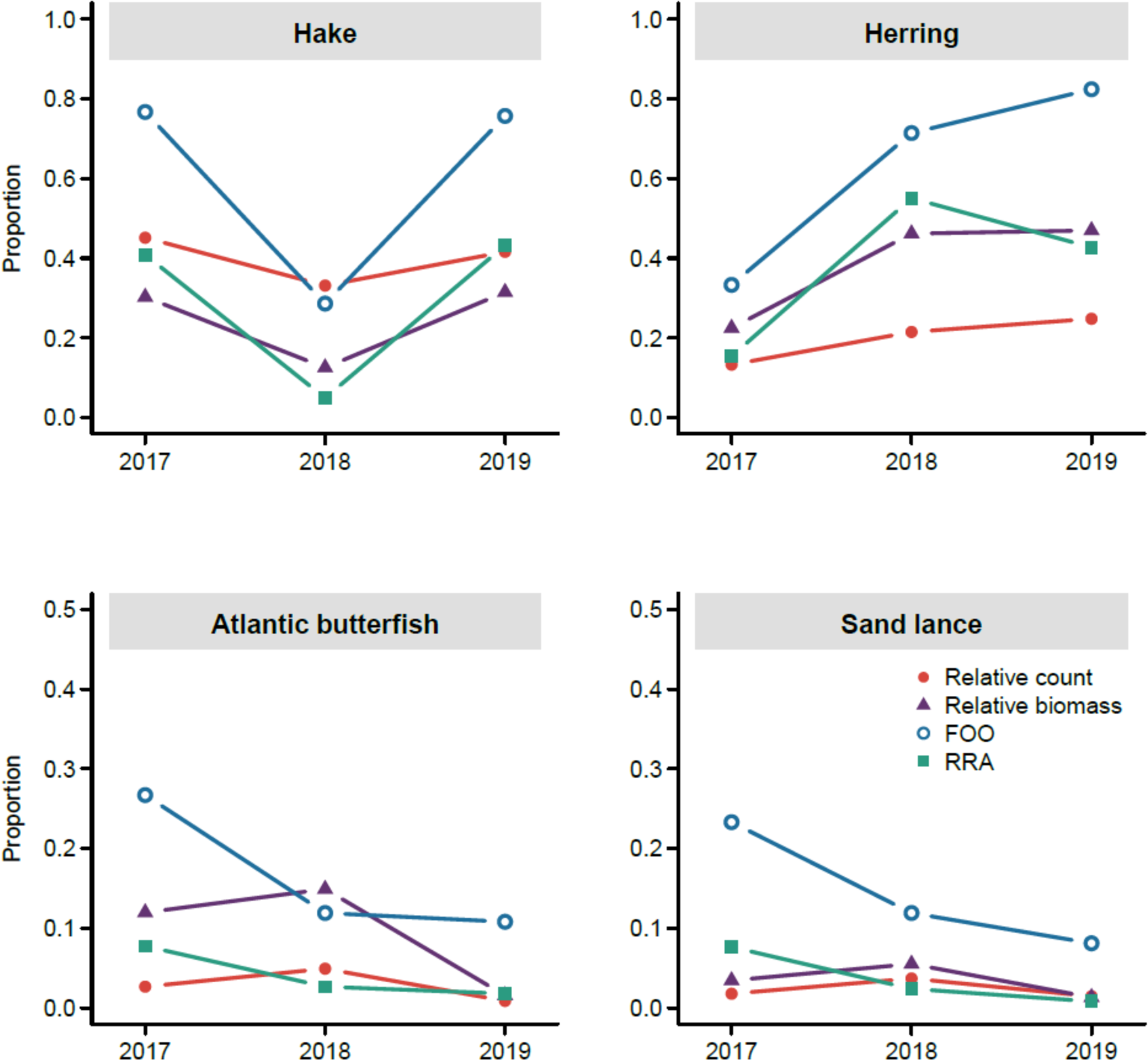
Interannual changes in diet metrics based on visual observations (relative counts, relative biomass) and dietary DNA (RRA, FOO) show variation in temporal patterns of diet composition based on the method used to characterize diet.

### 3.4 The effect of sample size on proportional estimates of diet

By simulating fecal datasets with sampling designs ranging from 10–40 successfully sequenced samples per year, we found that mean simulation error between RRA and relative biomass (measured by both RMSE and MAE) approached a lower asymptote at 35–40 samples/year (Figure 4). At a sample size of 40 samples/year, RRA and relative biomass had a mean RMSE of 0.08 (95% CI = 0.07, 0.09), and a mean MAE of 0.06 (95% CI = 0.05, 0.07). The interannual correlation between simulated RRA and relative biomass continued to increase between 10–40 samples/year ranging from a low of 0.28 (95% CI = -0.43, 0.80) at 10 samples/year to a maximum of 0.75 (95% CI = 0.72, 0.80) at 40 samples/year. This suggests that future studies should aim to collect at least 40 samples per year to accurately estimate diet proportions in this study system.

**Figure 4.**
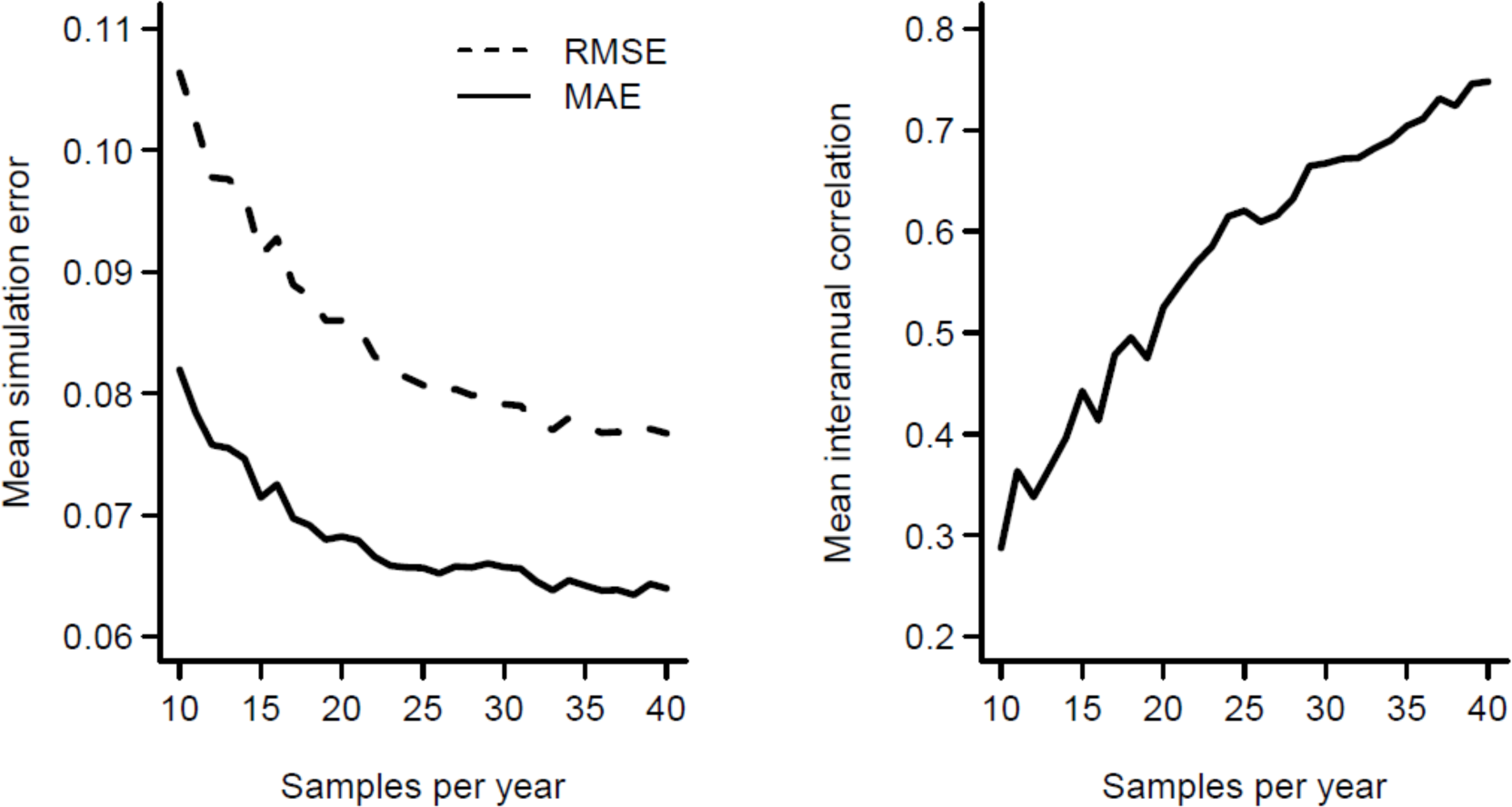
The match between diet metrics summarized from dietary DNA (RRA) and visual observations (relative biomass) increases with increasing fecal sample size. Error rates (mean absolute error [MAE] and root mean squared error [RMSE]) reach a lower asymptote and interannual correlations remain high at 35–40 samples per year. Error rates and correlations were generated from an iterative, simulated dietary DNA sampling scheme ranging from 10 to 40 samples per year compared against observed relative biomass.

## 4. Discussion

We used dietary DNA and visual observations of feeding events in a wild seabird population to determine to what extent dietary DNA could be used to estimate prey proportions. We found that the relative read abundance (RRA) of fish taxa recovered from fecal samples accurately captured the relative biomass consumed at the population-level over the three years of the study, with the relationship between RRA and relative biomass falling close to 1:1. The frequency of occurrence (FOO) of prey taxa, a commonly-used metric in dietary DNA studies, also showed a high correlation with relative biomass but over-estimated the importance of most fish taxa. In addition, we found that RRA slightly outperformed FOO in its ability to capture interannual changes in diet, and RRA produced similar estimates to relative biomass from high-effort visual observations. By simulating an annual fecal sampling program, we found that 35–40 successfully analyzed samples per year was necessary to produce RRA estimates that matched biomass estimates from visual observations. This range of sample effort provides a helpful benchmark for future dietary DNA studies aiming to characterize seabird diets and track climate- or human-induced change over time, although we caution that seabirds with more diverse diets (e.g., in the tropics) may require larger sample sizes.

Our study adds to the body of evidence that supports the use of RRA in dietary DNA studies (Deagle et al., 2019), particularly in avian systems (Verkuil et al., 2022). Deagle et al. (2019) suggested that RRA is a semi-quantitative metric, but encouraged its use over presence/absence metrics, such as FOO, because occurrence-based metrics can over-estimate the importance of prey consumed in small amounts (and low-level contamination). In our study, the very high correlation between RRA and relative biomass suggests that RRA is more than a semi-quantitative metric; it can give quantitative estimates of prey proportions in piscivorous seabirds. However, we caution that the low taxonomic resolution of visual observations in this study limited the number of prey categories that we could compare against dietary DNA metrics. In systems where the preyscape of seabirds is more diverse (or where prey can be compared at higher taxonomic resolution than was possible here), we may expect to see a larger benefit to using RRA over FOO due to the inflated impact of rare taxa on occurrence-based metrics. In other systems with especially high diet diversity, such as insectivorous birds or herbivorous mammals, validation studies and the use of mock communities are still advised.

Compared to traditional diet monitoring techniques, dietary DNA has some important advantages, particularly if we can use it to estimate prey proportions reliably as exhibited in this field test. Firstly, dietary DNA can provide greater taxonomic resolution than techniques that rely on the morphological identification of prey through either visual observations or the identification of hard parts, for example, from prey remains in fecal matter, regurgitates, or stomach contents (Hoenig et al., 2022). Expertise is often required to morphologically differentiate prey, and even then, species-level identifications can remain impossible. Observational studies can also suffer from observer bias and errors (Farmer et al., 2012), which are largely avoided with dietary DNA. In addition, a high number (almost one third) of our visual observations could not be identified at all due to the size, speed, or angle at which the fish were provisioned to chicks, whereas dietary DNA does not generally suffer from this limitation provided reference databases are complete. Dietary DNA can also provide greater taxonomic resolution than stable isotope-based methods, although, unlike stable isotope analyses, dietary DNA only provides a snapshot of the diet, while stable isotopes can integrate over longer time scales depending on the tissues analyzed (reviewed in Wiley et al., 2017). Dietary DNA also provides a means to collect diet information from a broader range of individuals with limited disturbance, for example from both adults and nestlings (Bowser et al., 2013; Fayet et al., 2021). This ability to rapidly ‘scale up’ population-level diet monitoring opens doors to more effectively track changes in prey availability over time and the potential impacts of these changes on sensitive populations (Barrett et al., 2007).

Although dietary DNA can provide complementary information on diet composition, it cannot estimate feeding rates, prey sizes, or prey quality. These metrics can have a significant effect on breeding success in seabirds (Davoren and Montevecchi, 2003; Fayet et al., 2021; Lamb et al., 2017; Schrimpf et al., 2012; Wanless et al., 2005) and often provide important information to guide seabird conservation efforts. Therefore, where existing diet monitoring programs exist, dietary DNA should be considered as a tool to increase the taxonomic resolution of the data collected and provide efficient estimates of diet composition as a supplement to field-based efforts. However, dietary DNA may prove especially useful when diet monitoring is constrained due to logistical challenges, such as an inability to observe feeding events or support observers in the field over extended periods of time. Since many seabird colonies are located on remote islands far from field stations, the ability to collect dietary DNA data during relatively short visits to colonies is a major advantage.

Overall, dietary DNA, when applied to seabirds, opens doors to more efficient, cost-effective tracking of climate- and fisheries-induced impacts on marine ecosystems. Dietary DNA has already been used to identify albatross populations with a heavy reliance on fishery discards, putting them at greater risk of bycatch (McInnes et al., 2017), and to elucidate the diets of endangered species such as the yellow-eyed penguin (Young et al., 2020). With the majority of fisheries resources already over-fished (Pauly et al., 1998) and climate-induced shifts in forage fish distributions (Kleisner et al., 2017; Pinsky et al., 2013), it is likely that seabird diets are shifting at unprecedented rates, yet we have not, until recently, had the tools to monitor this change in more than a few species. In some systems, estimates of forage fish availability may be more readily obtained from seabird diet data than through traditional net-based surveys (e.g. Scopel et al., 2017) and could prove especially valuable for numerous species and regions where data are limited or nonexistent (Cairns, 1988). Increasing the availability and accuracy of seabird diet data is timely, as fisheries management moves towards ecosystem approaches that aim to increase sustainability. By providing high-resolution estimates of prey proportions in seabird diets, we may provide the data necessary to parameterize food web models for ecosystem-based fisheries management (Hall and Mainprize, 2004; Pikitch et al., 2004) and track changes in forage fish availability on an annual basis using seabirds as indicators (Scopel et al., 2017; Velarde et al., 2013). Moving forward, the ability to monitor real-time changes in seabird diets through dietary DNA, combined with efforts to scale-up diet monitoring across populations, represents a key pathway to advance our mechanistic understanding of seabird declines and effectively address the challenges posed by fisheries management and global change.

## Acknowledgements

We would like to thank all the field and lab technicians that helped collect data for this project, including Elizabeth Ford, Taylor Ouelette, Caitlin Bowman, Amber Litterer, Aliya Caldwell, Olivia Smith, and Casey Coupe. Special thanks to Bronwyn Butcher, Irby Lovette, Jennifer Seavey, Adrienne Kovach, Brian Trevelline, and Jeffrey Hall for support and advice throughout. This study was supported by a seed grant from the University of New Hampshire School of Marine Science and Ocean Engineering. GC and AS were supported by postdoctoral fellowships from the Cornell Atkinson Center for Sustainability and Cornell Lab of Ornithology Rose Postdoctoral Program. The Isles of Shoals Tern Conservation Project is supported by the New Hampshire Fish and Game Department’s Nongame and Endangered Wildlife Program, the United States Fish and Wildlife Service (New Hampshire State Wildlife Grants), and the Shoals Marine Laboratory. We thank Michael Marchand and the New Hampshire Fish and Game Department, Nongame and Endangered Wildlife Program for leadership in New Hampshire tern conservation efforts. The handling of live birds was permitted under protocol #160403 approved by University of New Hampshire’s Institutional Animal Care and Use Committee and permit 06543 under the U.S. Geological Survey’s Bird Banding Laboratory. This paper is contribution 201 to the Shoals Marine Laboratory.

